# Neuroprotective effects of crocin I and II in an ischemia-reperfusion injury model

**DOI:** 10.1101/757971

**Authors:** Baowei Lv, Junyan Yin, Chunqing Feng, Yanhui Li

## Abstract

**Background:** Crocin I and II are derived from the medicinal plant *Crocus sativus* L. (Saffron), and their neuroprotective effects have been attracting more and more attention. However, their protective effect against cerebral apoplexy induced by hypoxia has not been reported. In this study, we aimed to clarify the roles of crocin I and II in protecting against ischemic injury.

**Materials/Methods:** We generated a rat cerebral ischemia-reperfusion injury model using a reversible cerebral artery occlusion suture method and found changes in amino acid neurotransmitters in the frontal cortex after drug administration. We also identified changes in mRNA expression of *Bcl2, Bax, Casp3, P38*, and *NFkb1* in the frontal cortex and changes in antioxidant indices in the brain.

**Results:** Crocin I and II both had protective effects on ischemic/anoxic injury *in vivo* by downregulating the expression of *Casp3* and *Nfkb1* mRNA and the steady-state levels of excitatory amino acids/inhibitory amino acids during ischemia and reperfusion and by improving the total antioxidant capacity and total superoxide dismutase activities during ischemia. We also found that crocin I and II had synergistic effects when used together.

**Conclusions:** These findings displayed that crocin I and II could protect animal model against ischemic and anoxic injury and provided new evidence for both molecules’ potential medicinal value.

## Background

Ischemic stroke is a primary cause of mortality and morbidity worldwide [1] and is now the leading cause of death in China [2]. However, there are limited options for the treatment of ischemic stroke [3,4], and only 2–5% of stroke patients receive thrombolytic therapy to restore blood flow because of the narrow post-stroke time window (<4.5 h) in which ischemic tissue can be rescued [5,6]. Recent advances in methods of treating ischemia-reperfusion, including neuroendovascular interventions, mechanical thrombectomy, in vivo imaging, stem cell therapy and computing programs, were developed.[7-10]. Animal studies have been performed to determine the pathological changes that occur after stroke and to explore the efficacy of new drugs and therapies.

The middle cerebral artery occlusion (MCAO) model in rats mimics the reperfusion of human ischemia after stroke and is used to study the pathological changes that occur after ischemic stroke and the mechanisms of drug action [11-13]. MCAO stroke induces cell death through interactions between excitatory amino acid (EAA) toxicity, acidosis, inflammatory responses, oxidative stress, peri-infarct depolarization, and apoptosis [14-16]. Both anoxia and ischemia increase the production of reactive oxygen species (ROS) [17-19] and induce inflammatory responses [19-21], and these cause damage to cellular components and negatively affect cell growth and function [22].

Crocin I and II (**Figure 1**) are purified from the medicinal plant *Crocus sativus* L. (Saffron) [23] and are widely reported to have antineoplastic properties [24-26]. Crocins were also found have the effect on metabolic syndrome-induced osteoporosis in rats [27]. Their neuroprotective effects have been attracting increasing attention; for example, the potential therapeutic effect of crocins for Alzheimer’s disease has been reported [23]. It has been also found that crocin could enhance hypothermia therapy in hypoxic ischemia-induced brain injury in mice [28]. The present study explored the efficacy and mechanisms of crocin I and II in the treatment of cerebral apoplexy induced by hypoxia in an MCAO model and investigated the neuroprotective effects of crocin I, crocin II, and a mixture of the two in *in vivo* models.

**Figure 1.**
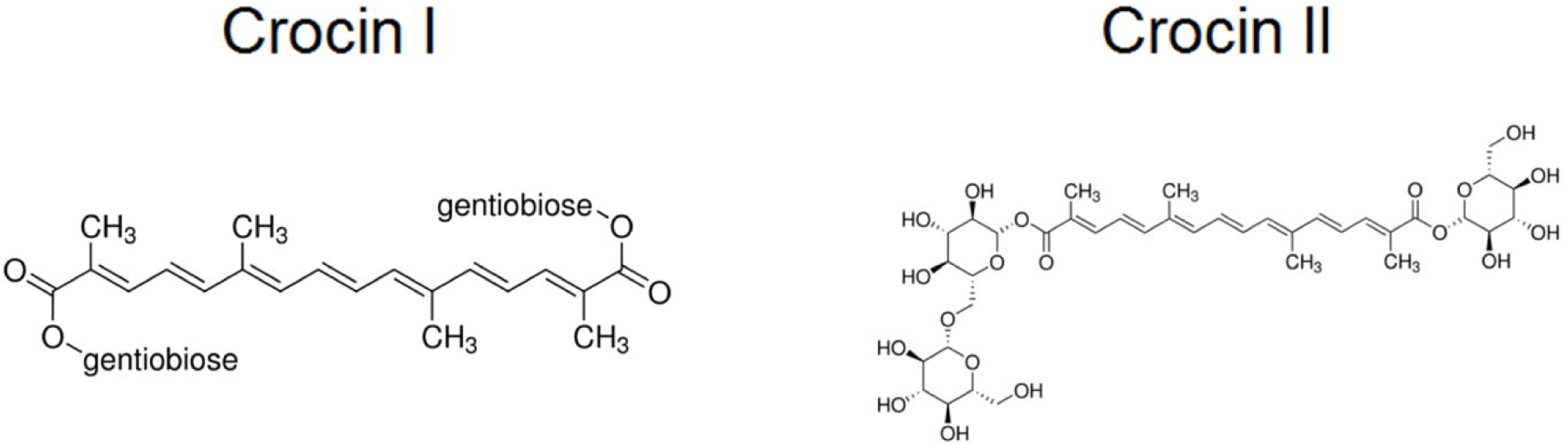
The structures of crocin I and II.

## Materials and methods

### Animals

Pathogen-free male SD rats (weight upon receipt: 120–140 g; weight at time of the experiments: 140–160 g) were provided by Beijing Vital River Laboratory Animal Technology Co., Ltd. (Beijing, China). The breeding environment consisted of a barrier-class animal laboratory [license number: SYXK(Lu)2013-0012] maintained at 20–25 °C with a 12:12 h light/dark cycle. The laboratory animals were cared for according to “*The Care and Use of Laboratory Animals*” by Liaocheng People’s Hospital.

### Rat cerebral ischemia-reperfusion injury model

The experimental groups included the sham group (Sham), MCAO-only group (Model), crocin I group (C1), crocin II group (C2), and crocin I and II (1:1) mixture group (C1+2). Both the Sham and Model groups were intragastrically administered 0.5% sodium carboxymethyl cellulose (Alibaba Co., Ltd., Hangzhou, China). The intragastrically administrated doses of crocin I (lot number: 111588-201303, purity: 92.6%), crocin II (lot number: 111589-201103, purity: 91.9%), and the crocin I and II mixture were 0.013 mg·kg^-1^ per day, 0.015 mg·kg^-1^ per day, and 0.014 mg·kg^-1^ per day, respectively, which were similar to the clinically relevant dose in humans. The intragastric administration volume was 10 mL·kg^-1^ and was given once per day for 14 days. After the 14-day treatments, rats from the Model, C1, C2, and C1+2 groups were anesthetized using 10% chloral hydrate at a dosage of 3.5 mL·kg^-1^. The MCAO operation was conducted using a reversible cerebral artery occlusion suture method that was designed for rats [29]. The Sham group underwent surgery similar to that of the MCAO group, but the external carotid artery and branches were not actually blocked.

### Amino acid sample preparation in the rat cortex

Six rats were allocated to each of the Sham, Model, C1, C2, and C1+2 groups as described above. At 40, 70, and 100 min after the MCAO operation and at 40, 100, and 200 min after reperfusion (referred to as I-40, I-70, I-100, I-100+R-40, I-100+R-100, and I-100+R-200, respectively), one rat in each group was decapitated and their brains were removed by quickly stripping the cerebral tissues, cutting off approximately 25 mg of frontal lobe tissue, and adding 1 mL of a methanol: water solution (volume ratio 1:1). The tissues were added to an ice bath with ultrasonication for 15 min and then homogenized by centrifugation at 13,000 × *g* for 30 s at 4°C. A portion of the supernatant (0.5 mL) was removed, and the solution containing the marker L-homoserine was added to reach a volume of 100 μg/L. After 15 min of centrifugation at 13,000 × *g* at 4°C, the supernatant was extracted for immediate use in the derivatization reaction.

### High-performance liquid chromatography

Glutamate (Glu) (lot number: 140690-201203, purity: 100%) were purchased from the National Institutes for Food and Drug Control (Beijing, China). Taurine (Tau) (lot number: T103829, purity: 99%), γ-aminobutyric acid (GABA) (lot number: A104200, purity: 99%), ethanethiol (lot number: E110411, purity: 98%), and o-phthalaldehyde (lot number: P108632, purity: 98%) were purchased from Shanghai Aladdin Biochemical Technology Co. Ltd. (Shanghai, China). High-performance liquid chromatography was performed on an Agilent 1260 (Agilent Technologies Singapore [International] Pte. Ltd., Singapore, Singapore) with precolumn derivatization. The software version used was ChemStation (Revision) B.04.03 (16). The derivatization reagent consisted of 27 mg of o-phthalaldehyde and 40 μL of ethanethiol dissolved in 5 mL of methanol to which 5 mL of 0.1 mmol/L sodium tetraborate buffer solution was added. Normal concentrations of amino acid solutions or supernatants from the previous section (40 μL) were added to 20 μL of the prepared derivatization solution. The solution was mixed for 30 s, allowed to stand for 120 s, and run through an Agilent ZORBAX C18 column (4.6 × 150 mm, 5 μm). Mobile phase A consisted of 10 mmol/L Na_2_HPO_4_:12H_2_O (pH 6.88), and mobile phase B consisted of methanol/acetonitrile (3:1). The ratio of phases A and B in solution was 6.4:3.6. The excitation wavelength was 355 nm, and the emission wavelength was 450 nm. The gain margin was ×6 with a lower sensitivity, the flow rate was set to 1.0 mL/min, the column temperature was 35°C, and the sample size was 10 μL.

### Antioxidant index determination

Ten rats each were allocated to the Sham, Model, C1, C2, and C1+2 groups as described above. In this experiment, the suture was taken out after 100 min of ischemia, and after reperfusion was performed for 20 h the rats were decapitated. The cerebral tissues were kept in cold normal saline to remove the blood and then were dried with filter paper. The left and right cerebral hemispheres were removed and weighed to prepare a 10% tissue homogenate. The kits used for detecting malondialdehyde (MDA) and glutathione peroxidase (GSH-Px) were purchased from Beijing Solarbio Science & Technology Co., Ltd. (Beijing, China) and Sigma-Aldrich Corporation (St. Louis, MO, USA) respectively. The kits used for detecting total antioxidant capacity (T-AOC) and superoxide dismutase (SOD) were both from Nanjing Jiancheng Bioengineering Institute (Nanjing, China). The relevant index detection was conducted according to the instructions of the kits described above. A UV-2550 ultraviolet spectrophotometer (Shimadzu Scientific Instruments, Kyoto, Japan) was used for detection.

### Real-time polymerase chain reaction

Twelve rats each were allocated to the Sham, Model, C1, C2, and C1+2 groups as described above. After the MCAO operation, one rat from each group was sacrificed by decapitation at I-40, I-100, I-100+R-40, and I-100+R-200. RNA Lyzol (Shanghai ExCell Biology, Inc., Shanghai, China) was used for RNA extraction. A RevertAid First Strand cDNA Synthesis Kit (Thermo Fisher Scientific Inc., Waltham, MA, USA) was used for reverse transcription, and 5 μg of RNA was used for each sample. RT-PCR Master Mix (Toyobo Co., Ltd., Osaka, Japan) was used on an ABI 7500 fast real-time fluorogenic quantitative RT-PCR system (Applied Biosystems, Foster City, CA, USA). The primers used in the real-time polymerase chain reaction were P-P38-F: 5’-GA ATG GAA GAG CCT-3’; P-P38-R: 5’-GAC AGA ACA GAA GC-3’; NF-κB-F: 5’-CGA CAC CTC TAC AC-3’; NF-κB-R: 5’-GGC TCA AAG TTC TC-3’; BCL2-F: 5’-A CAA CAT CGC TCT-3’; BCL2-R: 5’-CA GGA GAA ATC AAA-3’; BAX-F: 5’-CCA CCA AGA AGC-3’; BAX-R: 5’-A CGG AAG AAG ACC-3’; caspase-3-F: 5’-CTT GGA AAG CAT C-3’; caspase-3-R: 5’-AGC CTG GAG CAC AG-3’; actin-F: 5’-TC TAT GAG GGT TAC-3’; and actin-R: 5’-GTC ACG CAC GAT TTC-3’.

### Statistical analysis

All data are presented as the mean ± standard deviation and were analyzed with GraphPad Prism 6.01 statistical software (GraphPad Software, Inc., La Jolla, CA, US). The t-test was used for analyzing measurement data. Differences between two groups were analyzed by using the Student’s t-test. Comparison between multiple groups was done using One-way ANOVA test followed by Post Hoc Test (Least Significant Difference). *P*<0.05 was considered statistically significant.

## Results

### Effects of crocin I and II on cortical amino acids in the rats with MCAO-induced cerebral ischemia-reperfusion injury

Compared with the Sham group, the Glu concentrations in the Model group were significantly increased at each time point of cerebral ischemia and reperfusion (**Figure 2A**, *P*<0.05). Compared with the Model group, the Glu concentrations in the C1, C2, and C1+2 groups were decreased at the I-100 and I-100+R-40 time points, and in the C2 and C1+2 groups the Glu concentrations were decreased at the I-100+R-70 and I-100+R-200 time points (**Figure 2A**, *P*<0.05). We also found that Glu levels in the C1+2 group were lower than either the C1 or C2 groups at the I-100+R-40, I-100+R-70, and I-100+R-200 time points (**Figure 2A**, *P*<0.05), suggesting that administration of crocin I and II at the same time has a stronger effect than using only one alone.

**Figure 2.**
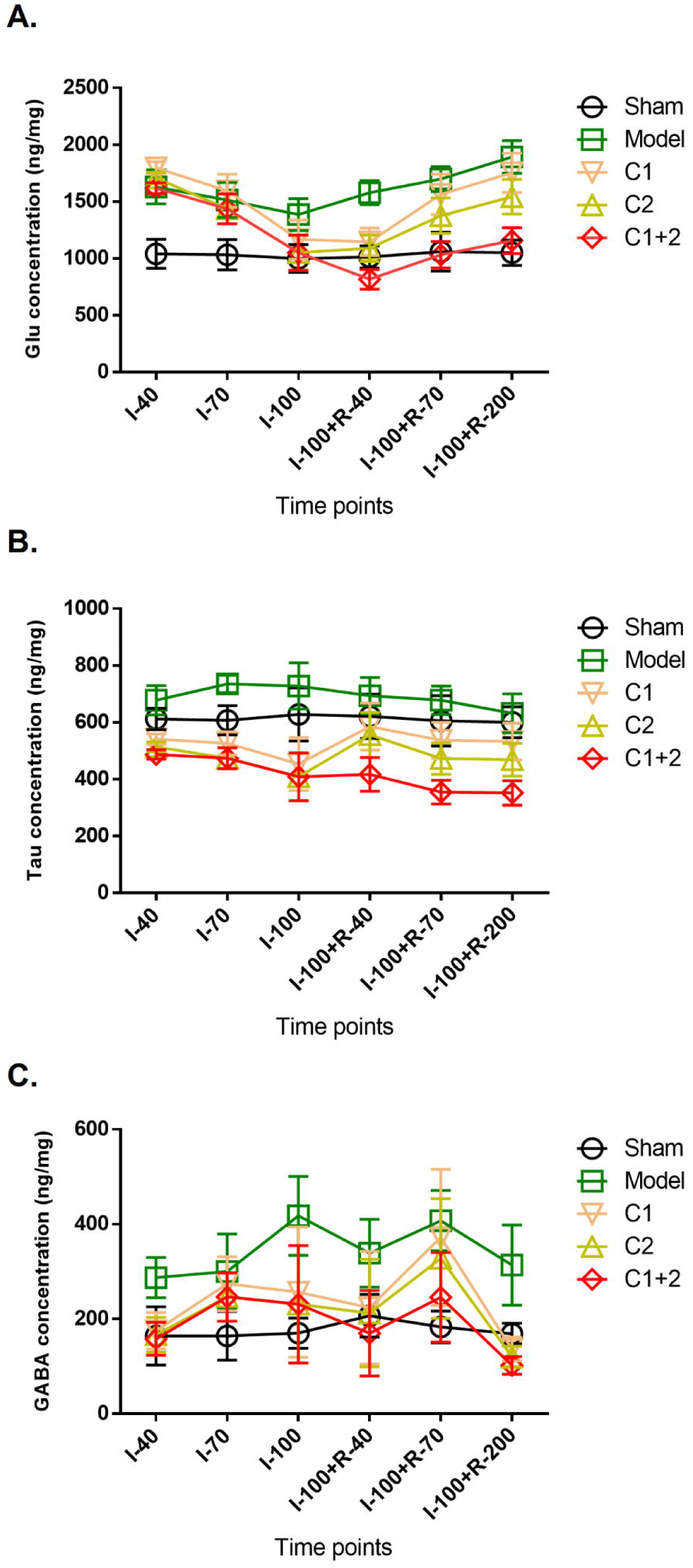
The changes in Glu (A), Tau (B), and GABA (C) concentrations in the frontal cortex in Model rats after crocin I and II intervention for ischemia and reperfusion (ng/mg, n = 6). I-40: ischemia for 40 min; I-70: ischemia for 70 min; I-100: ischemia for 100 min; I-100+R-40: ischemia for 100 min, then reperfusion for 40 min; I-100+R-70: ischemia for 100 min, then reperfusion for 70 min; I-100+R-200: ischemia for 100 min, then reperfusion for 200 min.

Compared with the Sham group, the Tau concentrations in the Model group were significantly increased at I-70 and I-100 (**Figure 2B**, *P*<0.05). Compared with the Model group, the Tau concentrations were decreased at all six time points in the C1, C2, and C1+2 groups (**Figure 2B**, *P*<0.05). Notably, the Tau level in the C1+2 group was lower than either the C1 or C2 groups at I-100+R-40, I-100+R-70, and I-100+R-200 (**Figure 2B**, *P*<0.05).

Compared with the Sham group, the GABA concentrations in the Model group were significantly increased at I-70, I-100, I-100+R-40, I-100+R-70, and I-100+R-200 (**Figure 2C**, *P*<0.05). Compared with the Model group, the GABA concentrations in at least one treatment group were decreased at I-40, I-100, I-100+R-40, I-100+R-70, and I-100+R-200 (**Figure 2C**, *P*<0.05). Notably, the GABA level in the C1+2 group was lower than the C1 group at I-100+R-70 (**Figure 2C**, *P*<0.05).

### Effects of crocin I and II on antioxidant indices in MCAO-induced cerebral ischemia-reperfusion injury

None of the four antioxidant indices were significantly changed in the left-brain tissue (the non-ischemic side) of the rats in the Model group or crocin-administered groups compared with the Sham group (**Figure 3**, *P*>0.05). MDA levels were significantly increased in the right-brain tissue (the ischemic side) in the Model group (**Figure 3A**, *P*<0.05), while GSH-Px, T-AOC, and total SOD (T-SOD) levels were significantly decreased (**Figure 3B-3D**, *P*<0.05). Compared with the Model group, the T-AOC and T-SOD activities were significantly increased in the right-brain tissue of the C1+2 group (**Figure 3B-3C**, *P*<0.05).

**Figure 3.**
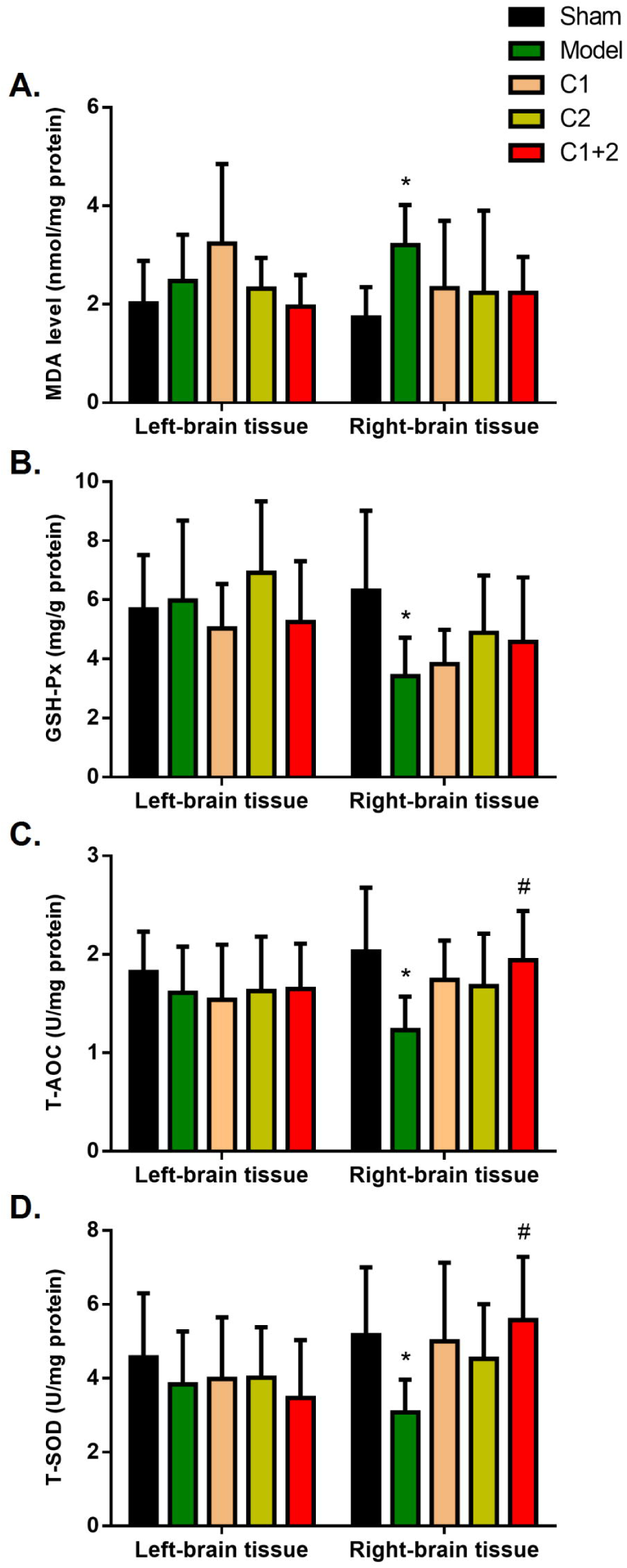
Antioxidant index results, including MDA (A), GSH-Px (B), T-AOC (C), and T-SOD (D), for the left-brain tissue (non-ischemic control side) and right-brain tissue (ischemic side). n = 10. *: *P*<0.05 *vs*. the Sham group; #: *P*<0.05 *vs*. the Model group.

### Effects of crocin I and II on p38, Nfkb1, Bcl2, Bax, and Casp3 mRNA expression in the frontal lobe of rats with MCAO-induced cerebral ischemia-reperfusion injury

The *Bcl2*/*Bax* ratio of mRNA expression was significantly decreased in the Model group compared with the Sham group at all four time points (**Figure 4A**, P<0.05). Compared with the Model group, the ratio was not upregulated in the C1, C2, or C1+2 groups (**Figure 4A**, *P*>0.05). The *Casp3* and *Nfkb1* mRNA expression levels were significantly higher in the Model group compared to the Sham group (**Figure 4B-4C**, *P*<0.05), and they were downregulated in the C1, C2, and C1+2 groups at all four time points (**Figure 4B-4C**, *P*<0.05). At I-100+R-200, the *Casp3* mRNA level in the C1+2 group was significantly lower than in the C1 group, while there were no differences between the C1 and C2 groups (**Figure 4C**, *P*<0.05). The *Nfkb1* mRNA level in the C1+2 group was lower than in the C1 and C2 groups (**Figure 4C**, *P*<0.05), and the *p38* mRNA expression levels were unchanged among groups (**Figure 4D**, *P*>0.05).

**Figure 4.**
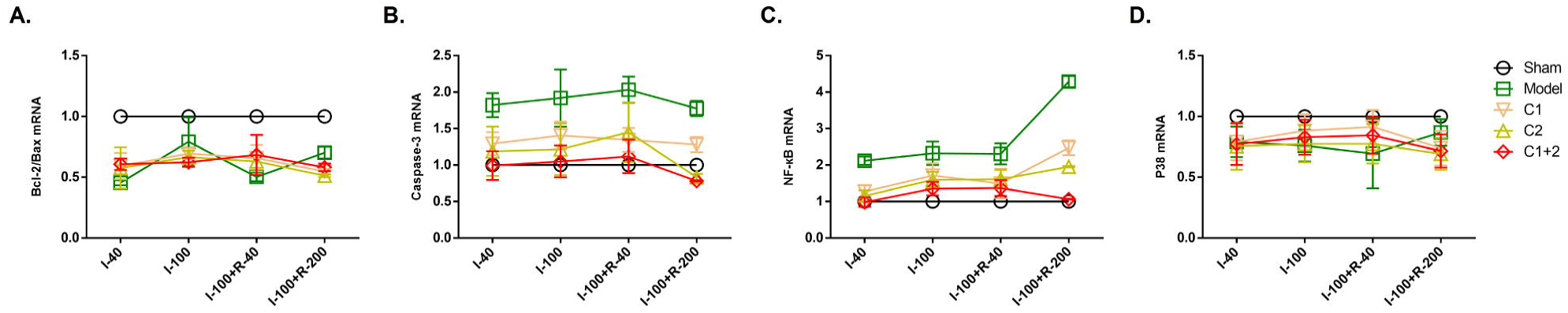
Effects of crocin I and II on the mRNA levels of the *Bcl2*/*Bax* ratio (A), *Casp3* (B), *Nfkb1* (C), and *p38* (D) in the prefrontal cortex of ischemia-reperfusion injury rats. n = 3. I-40: ischemia for 40 min; I-100: ischemia for 100 min; I-100+R-40: ischemia for 100 min, then reperfusion for 40 min; I-100+R-200: ischemia for 100 min, then reperfusion for 200 min.

## Discussion

We generated a rat cerebral ischemia-reperfusion injury model using a reversible cerebral artery occlusion suture method. Since crocins were used as preventative treatment for ischemia-reperfusion injury, they were actually administrated before ischemia-reperfusion injury modelling. We then further discussed the pathways involving inflammatory factors (p38 and NF-κB), apoptotic factors (Bcl-2, Bax, and caspase-3), amino acid neurotransmitters, and ROS that contribute to learning and memory in the cortex with drug intervention in order to elucidate the effects and properties of crocin I and II and thereby provided support for their clinical applications.

The neurotoxic properties of EAAs result in the initiation and development of cerebral tissue injury, and Glu is one of the most important EAAs in the brain [30]. Tau and GABA are important inhibitory amino acids (IAAs) in the brain, and in models of cerebral ischemia they are released by cells to counteract the effects of EAAs [31,32]. The present study was conducted to assess the neuroprotective mechanisms of crocin I and II by measuring changes in amino acid concentrations in the cerebral cortex at different time points after ischemia-reperfusion injury. The Glu/GABA ratio is a measure of the balance of cerebral EAAs and IAAs, and we found that in ischemia-reperfusion injury the *in vivo* Glu and GABA levels were increased (**Figure 2A** and **2C**) as were the corresponding overall EAA/IAA ratios. In all three crocin-treatment groups, the concentrations of certain amino acids that are involved in EAA/IAA homeostasis were downregulated at different time points and they had a tendency to approach normal levels. Thus crocin I and II appear to play a protective role in the acute stage of ischemia-reperfusion by reducing the neurotoxicity of EAAs. Crocin I and II decreased the concentrations of the IAAs Tau and GABA (**Figure 2B-2C**), and the Glu/GABA ratio was maintained at normal homeostatic levels and thus reduced the toxicity of EAAs. Maintaining and improving the relative concentrations of EAA and IAA might be one of the mechanisms underlying crocin I and II’s action in reducing the cerebral infarction volume in acute cerebral ischemia.

The T-AOC of an organism reflects its ability to resist oxidation and to scavenge free radicals [33] and consists of enzymatic and nonenzymatic antioxidant defense systems that include, but are not limited to, SOD, catalase, GSH-Px, vitamin C, vitamin E, glutathione, glucose, and β-carotene. The T-AOC is more dependable for evaluating whether stress has caused oxidative damage to an organism than information provided by a single antioxidant index. When endogenous or exogenous events cause abnormalities in an organism’s metabolism that lead to the sudden production of large amounts of ROS, the antioxidant defense system will be triggered and excessive ROS will be removed, thereby protecting the tissues from oxidative damage [34]. SOD is an endogenous antioxidant enzyme that metabolizes ROS into hypotoxic substances, hence protecting cells from damage and playing a crucial role in balancing oxidation and antioxidation in an organism [35,36]. Nonenzymatic ROS attacks on polyunsaturated fatty acids in biological membranes form lipid peroxides such as MDA, and thus the MDA concentration reflects the degree of lipid peroxidation and indirectly reflects the degree of cell damage. The current study also examined the antioxidation mechanisms of crocin I and II in protecting against ischemic cerebral injury and their influence on antioxidant indices in bilateral ischemic and non-ischemic rat cerebral tissues. Crocin I and II significantly improved T-SOD and T-AOC activity in bilateral ischemic cerebral tissues (**Figure 3C-3D**), suggesting that *in vivo* antioxidant enzyme synthesis rescues the aberrant oxidation/antioxidation balance seen in oxidative-stress injury and that crocin I and II play important roles in the enzymatic and nonenzymatic antioxidant defense systems to protect cells from oxidation damage.

We also found that when crocin I and II were administered prior to MCAO-induced ischemia-reperfusion injury in rats, the expression of *Casp3* and *Nfkb1* mRNA was downregulated at different time points after ischemia reperfusion (**Figure 4B-4C**). These results suggest that crocin I and II play a protective role in ischemia-reperfusion injury through the downregulation of factors associated with apoptosis.

Interestingly, we also found that using crocin I and II together with both at half dosages had better effects at some time points than administering only crocin I or crocin II alone at the full dosage (**Figure 2-4**), thus suggesting that crocin I and II have synergistic neuroprotective effects against *in vivo* ischemic and anoxic injury.

To be honest, we also compared infract volumes between C1+2 and Model groups using TTC staining and *in vivo* MR imaging. Results exhibited no statistical significance (not shown) between groups for both methods. As crocins were designed as preventative treatment for ischemia-reperfusion injury in current study, they were administrated prior to modeling. Though crocins could not significantly reduce the infract volume caused by ischemia-reperfusion injury, they restored levels of important molecular makers of ischemia-reperfusion injury including amino acid neurotransmitters, antioxidant indexes, inflammatory factors and key molecules in apoptosis signaling.

## Conclusions

We found that crocin I and II exerted protective effects against *in vivo* ischemic and anoxic injury and exhibited synergistic effects when used together. This research contributes to a greater understanding of the mechanistic functions of crocin I and II. We will further evaluate the neural functioning, including behaviors such as modified neurological severity score (mNSS) and foot-fault degree, permeability of the blood brain barrier BBB, *in vivo* morphology of brain and neurobiological marker staining for tissues of the infarction and surrounding area, to determine whether crocin protection improves prognosis in the future work.

